# Mutational signatures reveal the role of RAD52 in p53-independent p21 driven genomic instability

**DOI:** 10.1101/195263

**Authors:** Panagiotis Galanos, George Pappas, Alexander Polyzos, Athanassios Kotsinas, Ioanna Svolaki, Nikos N Giakoumakis, Christina Glytsou, Ioannis S Pateras, Umakanta Swain, Vassilis Souliotis, Alexander Georgakilas, Nicholas Geacintov, Luca Scorrano, Claudia Lukas, Jiri Lukas, Zvi Livneh, Zoi Lygerou, Claus Storgaard Sørensen, Jiri Bartek, Vassilis G. Gorgoulis

**Affiliations:** Molecular Carcinogenesis Group, Department of Histology and Embryology, School of Medicine, National Kapodistrian University of Athens, 75 Mikras Asias Str, Athens, GR-11527, Greece.; Danish Cancer Society Research Centre, Strandboulevarden 49, Copenhagen, DK-2100, Denmark.; Biomedical Research Foundation of the Academy of Athens, 4 Soranou Ephessiou Str, Athens, GR-11527, Greece.; Laboratory of Biology, School of Medicine, University of Patras, 26505, Rio, Patras, Greece; Department of Biology, University of Padova, Padova 35121, Italy; Dept. of Biomolecular Sciences, Weizmann Institute of Science, Rehovot, 76100, Israel; Institute of Biology, Medicinal Chemistry and Biotechnology, National Hellenic Research Foundation, 48 Vassileos Constantinou Ave, Athens, GR-11635, Greece; Physics Department, School of Applied Mathematical and Physical Sciences, National Technical University of Athens (NTUA), Zografou 15780, Athens, Greece; Dept. of Chemistry, New York University, New York 10012, USA; Novo Nordisk Foundation Center for Protein Research, Faculty of Health and Medical Sciences, University of Copenhagen, Copenhagen, Denmark; Biotech Research and Innovation Centre (BRIC), University of Copenhagen, Ole Maaloes Vej 5, Copenhagen, DK-2200, Denmark; Institute of Molecular and Translational Medicine, Faculty of Medicine and Dentistry, Palacky University, Hněvotínská, Olomouc, 1333/5 779 00, Czech Republic.; Science for Life Laboratory, Division of Genome Biology, Department of Medical Biochemistry and Biophysics, Karolinska Institute, Stockholm, SE-171 77, Sweden.; Faculty of Biology, Medicine and Health, University of Manchester, Manchester Academic Health Science Centre, Wilmslow Road, Manchester, M20 4QL, UK

**Keywords:** p21^WAF1/Cip1^, Rad52, Genomic Instability, Translesion DNA Synthesis-TLS, Single Nucleotide Substitution (SNS), Break Induced Replication-BIR, Single Strand Annealing-SSA

## Abstract

**Background:** Genomic instability promotes evolution and heterogeneity of tumors. Unraveling its mechanistic basis is essential to design appropriate therapeutic strategies. In a recent study we reported an unexpected oncogenic property of p21^WAF1/Cip1^ showing that its chronic expression, in a p53-deficient environment, causes genomic instability by deregulating the replication licensing machinery.

**Results:** Extending on this work we now demonstrate that p21^WAF1/Cip1^ can further fuel genomic instability by suppressing the repair capacity of low and high fidelity pathways that deal with nucleotide abnormalities. Consequently, fewer single nucleotide substitutions (SNSs) occur, while formation of highly deleterious DNA double-strand breaks (DSBs) is enhanced, crafting a characteristic mutational signature landscape. Guided by the mutational signatures formed, we found at the mechanistic level that the DSBs were repaired by Rad52-dependent Break-Induced Replication (BIR) and Single-Strand Annealing (SSA). Conversely, the error-free synthesis-dependent strand annealing (SDSA) repair route was deficient. Surprisingly, Rad52 was activated transcriptionally in an E2F1-dependent manner, rather than post-translationally as is common for DNA repair factor activation.

**Conclusions:** Our results signify the importance of mutational signatures as guides to disclose the “repair history” leading to genomic instability. In this vein, following this approach we unveiled how chronic p21^WAF1/Cip1^ expression rewires the repair process, identifying Rad52 as a source of genomic instability and a candidate therapeutic target.

## Background

Genomic instability is a hallmark of cancer that plays an important role in shaping tumor behavior over time **[1-5]**. Elucidating the molecular routes that drive this phenomenon is essential for designing proper therapeutic strategies as well as monitoring the natural history of carcinogenesis **[6]**. Not all genetic alterations are *driver* events, since each type of cancer bears a large number of *passenger* mutations **[7].** Although the latter have no causative role in cancer development they represent pieces of a *mutational signature* pattern that can provide information about the type(s) of DNA damage taking place and the repair pathway(s) involved **[7, 8].**

We have recently reported that precancerous- and cancerous-associated chronic p21^WAF1/Cip1^ expression, in a p53-deficient environment, fuels genomic instability by deregulating the replication licensing machinery causing re-replication, a deleterious form of replication stress. These events occur throughout a senescence-like phase during which an error-prone DNA repair process takes place forming a genetic landscape that allowed a subpopulation of p21^WAF1/Cip1^ – expressing cells to escape senescence (termed “escape cells”) **[9]** (**Fig. 1a)**. Interestingly and in accordance with the oncogene-induced DNA damage model for cancer development **[4]**, the “escaped cells” demonstrated aggressive features and increased chemo-resistance **[9]** (**Fig. 1a)**.

**Figure 1.**
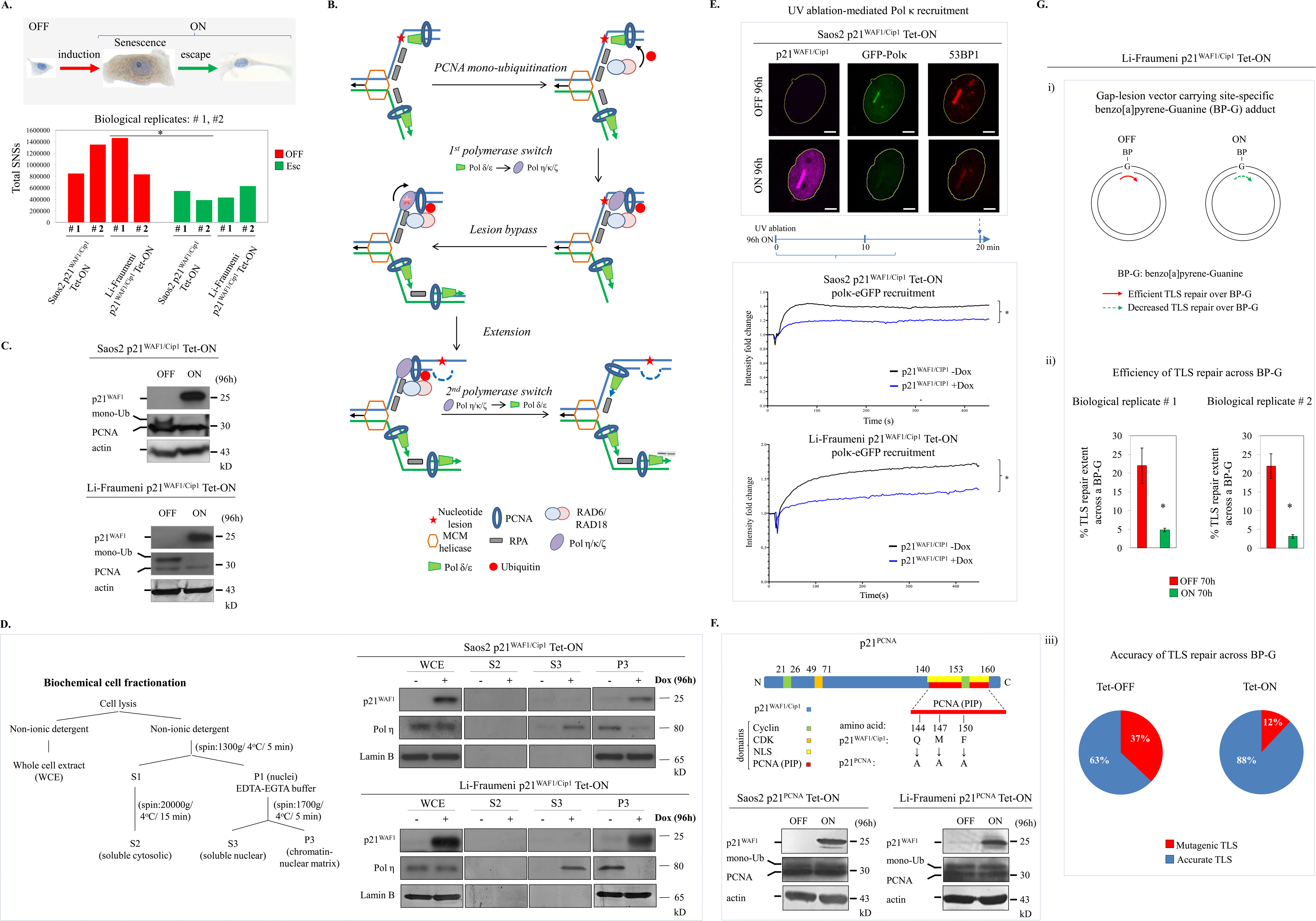
Reduction of Single Nucleotide Substitution (SNSs) and malfunction of the translesion DNA synthesis and repair (TLS) process upon protracted p21^WAF1/Cip1^ expression. (**a**) Chronic p21^WAF1/Cip1^ expression, in a p53-deficient environment, leads to the emergence of a subpopulation of p21^WAF1/Cip1^ aggressive and chemo-resistant (“escaped”: Esc) cells (right-top photo depicting escaped Li-Fraumeni-p21^WAF1/Cip1^ Tet-ON cell) after bypassing an initial senescence-like phase (middle-top photo depicting senescent Li-Fraumeni-p21^WAF1/Cip1^ Tet-ON cell stained with a new senescence detecting compound [**70**]) that carry a lower SNS “load” depicted in histograms (below photos)) [* *p*<0.05 (Saos2), * *p*=0.05 (Li-Fraumeni“OFF” vs “ESCAPED”), Welch’s t-test, error bars indicate SDs, n=2 biological replicates]. (**b**) Scheme of TLS pathway function. TLS is a DNA damage tolerance process enabling the DNA replication machinery to replicate over DNA lesions. Upon DNA damage PCNA is mono-ubiquitinated, followed by polymerase switch from normal high-fidelity DNA replication polymerases to TLS ones. TLS polymerase Polη, bound to PCNA, inserts a nucleotide opposite to the lesion and assisted by an additional TLS polymerase like Polκ or Polζ, extends beyond the insertion. Finally, a second polymerase switch takes place by substituting TLS polymerases with high-fidelity ones. (**c**) Sustained p21^WAF1/Cip1^ expression results in decreased mono-ubiquitination of PCNA (mono-Ub). Immunoblots (IBs) in 96h induced Saos2- and Li-Fraumeni-p21^WAF1/Cip1^ Tet-ON cells. (**d**) Reduced binding of Pol η to chromatin in cells with protracted p21^WAF1/Cip1^ expression. IBs after cell fractionation (described in scheme) depicting lower levels of Pol η in chromatin extracts from 96h induced Saos2- and Li-Fraumeni-p21^WAF1/Cip1^ Tet-ON cells. (**e**) Immunofluorescent confocal microscopy (top panel) showing reduced Polκ loading on regions of damaged chromatin after UV-laser ablation in 96h induced Saos2-p21^WAF1/Cip1^ Tet-ON cells transfected with a GFP-Polκ vector. Plots (lower panel) depict recruitment kinetics of Polκ in Saos2- and Li-Fraumeni-p21^WAF1/Cip1^ Tet-ON cells, respectively (see also **Suppl Video 1a-d**). The average intensity of fluorescence at the site of damage and the total cell fluorescence in respect to time were quantified and plotted. Five cells in each condition of two independent experiments were processed. Time frames for obtaining IFs and recruitment plots are depicted in middle scheme. (**f**) Scheme describes a specific p21^WAF1/Cip1^ mutant [p21^PCNA^: harboring Q144, M147, F150 substitutions to A in its PCNA-interacting-protein (PIP) degron motif] abrogating its interaction with PCNA [**9**]. IBs depict mono-ubiquitination of PCNA (mono-Ub) in 96h induced Saos2- and Li-Fraumeni-p21^PCNA^ Tet-ON cells. (**g**) Overexpression of p21^WAF1/Cip1^ decreases TLS repair efficiency across a site-specific lesion in a gapped plasmid TLS assay (i). Induced Li-Fraumeni-p21^WAF1/Cip1^ Tet-ON cells were assayed for TLS efficiency (ii) and accuracy of repair (iii) with a gap-lesion vector carrying a site-specific benzo[a]pyrene-Guanine (BP-G) adduct (i) (see also **Suppl Table 1**). Actin and Lamin B serve as loading control.

Which particular error-prone repair pathway(s) is(are) employed by the p21^WAF1/Cip1^ expressing cells to craft the permissive environment for “senescence escape” is a key question, as its answer would unveil potential genomic instability routes that could represent future therapeutic targets. To address this question we followed a reverse engineering approach examining the *mutational signature* patterns of the “escaped cells”. The *mutational signatures* that include amount and type of single nucleotide substitutions (SNSs), insertions, deletions (INDELs) and breakpoint junctions reflect the actual “repair history” that takes place following exogenous and/or endogenous mutagenic events. As a guideline we utilized the 21 distinct *mutational signatures*, reported by Alexandrov and collegues, who extracted them analysing ∼5×10^6^mutations from ∼7000 cancers [**8**]. We demonstrate that p53-independent elevated p21^WAF1/Cip1^ expression drives genomic instability by rewiring the global cellular DNA repair landscape towards predominantly error-prone processes that prominently rely on the RAD52 recombinase.

## Results

### 1. The Single Nucleotide Substitution (SNS) load is reduced in p21^WAF1/Cip1^ escaped cells

As a first step we evaluated the single-nucleotide-substitution (SNS) load in the human p21^WAF1/Cip1^ escaped cell models **[9]** and found that they harbor fewer SNSs compared to the p21^WAF1/Cip1^ -un-induced controls **(Fig. 1a).** This result was cell-type independent, consistently observed in both p21^WAF1/Cip1^ -inducible cellular systems examined, namely cancerous Saos2 cells and non-cancerous cells from a Li-Fraumeni syndrome patient **(Fig. 1a).** Given that protracted p21^WAF1/Cip1^ expression in a p53-null environment promotes genomic instability **[9]**, the reduced number of SNSs in the escaped cells seemed at first glance counter-intuitive. Yet, from a broader perspective, even though SNSs represent genetic defects, they signify the “last option” the cell possesses to avoid the more detrimental double strand break (DSB) lesions **[7]**. TLS or DNA damage tolerance (DDT) is the main cellular repair mode that orchestrates the choice of the above mentioned “last option” by dealing with all types of SNSs that remain unrepaired by the cellular high fidelity repair mechanisms. TLS is also known as post replication repair (PRR) as it initiates DNA synthesis downstream of a DNA lesion, thus allowing repair after DNA synthesis **(Fig. 1b) [10,11].** Hence, the reduced amount of SNSs in the “escaped” cells could result from a malfunctioning TLS/DDT repair process and/or declined activity of one or more high fidelity repair pathways that commonly mend defective nucleotides.

### 2. Reduced SNS load reflects malfunctioning TLS and excision repair pathways BER and NER

A key step regulating TLS is monoubiqutylation of PCNA (Ub-PCNA), at Lys 164 (K164), carried out by the E3-ligase Rad18 **[12]**. Such monoubiquitylated PCNA (Mono-Ub-PCNA) operates as a “molecular switch” shifting normal DNA replication into TLS. Mono-Ub-PCNA increases its affinity for TLS polymerase Polη, a Y-family polymerase that inserts a nucleotide directly opposite of a nucleotide lesion **(Fig. 1b) [13].** Although Polη is involved in error-free lesion bypass of UV-induced thymidine (TT) dimers **[14, 15]**, it can also catalyse error-prone bypass of other lesion types such as 8-oxoguanine, apurinic/apyrimidinic (AP) sites and DNA adducts caused by benzo[a]pyrene diol epoxide (BPDE), leading to mutagenic consequences **[16].** Following p21^WAF1/Cip1^ induction both, the mono-Ub-PCNA and the chromatin bound fraction of Polη were dramatically reduced, consistent with our conjecture that dysfunctional TLS may be responsible for the reduced SNS load in the escaped cells **(Fig. 1c, d).** In line with the above, recruitment of Polκ, an extender in translesion synthesis [**17**], to sites of UV-laser induced DNA damage was significantly decreased in cells expressing p21^WAF1/Cip1^ (**Fig. 1e; Suppl Video 1a-d)**. Supporting the notion that the induced p21^WAF1/Cip1^ binds to and inhibits PCNA monoubiqutylation, and consequently reduces recruitment of the TLS polymerases **[18],** replacing the wild-type p21^WAF1/Cip1^ by a p21 mutant defective in PCNA binding (p21^PCNA^) did not reduce PCNA monoubiqutylation (**Fig. 1f)**. To assess whether the above biochemical traits reflect dysfunctional TLS repair we employed a gapped plasmid TLS assay [**19**] to quantify the extent of repair across a site-specific (benzo[a]pyrene-guanine) adduct and found that induction of p21^WAF1/Cip1^resulted in a robustly decreased (4.6-7.1 fold) frequency and concomitantly lower repair accuracy (12% instead 37%) **(Fig. 1g, Suppl Table 1)**.

An additional feature that could further exacerbate the impact of malfunctioning TLS-mediated repair would be a potentially increased eroneous nucleotides in the genome due to deregulation of the high-fidelity nucleotide repair mechanisms. It has been reported that p21^WAF1/Cip1^can negatively modulate high fidelity DNA repair processes, particularly those implicated in excising defective nucleotides, such as nucleotide excision repair (NER), mismatch repair (MMR) and base excision repair (BER) **[20].** Given that our experimental systems were not exposed to exogenous causes of DNA damage, such as UV or X-ray irradiation, a major potential source of nucleotide abnormalities could be an endogenous process, particularly over-production of reactive oxygen species (ROS) [**21**]. Elevated ROS would lead to nucleotide oxidative lesions, with 8-oxo-dGuanine (8-oxo-dG) being the most frequent one **[22].** Indeed, p21^WAF1/Cip1^ induction was followed by a progressive generation of ROS **(Fig. 2a),** in agreement with previous findings **[23]**. Applying a modified alkaline Comet assay we could detect 8-oxo-dG in the DNA of the p21^WAF1/Cip1^ -induced cells using OOG1 (8-Oxoguanine glycosylase) (**Fig. 2c**). Moreover, RNAseq analysis showed that essential factors of the nucleotide repair mechanisms were down-regulated **(Fig. 2b, 3a).** Particularly, BER, which is mainly responsible for removing oxidative lesions **[22]** was affected, as the levels of several key apical glycosylases and downstream effectors were down-regulated **(Fig. 2b, Suppl. Fig 1).** Consistently, we found enhanced DNA incorporation of 8-oxo-dG in p21^WAF1/Cip1^ expressing cells, using an 8-oxo-dG specific assay [**24**], indicative of a lower OGG1 activity **(Fig. 2c).** NER was disrupted as well, as judged from the levels of its components and the repair capacity of N-alkylpurine monoaducts **[25, 26] (Fig. 3b, Suppl. Fig 1a,c).** Although NER is primarily involved in repairing bulky DNA lesions, for instance UV-induced TT dimers, it can also repair non-bulky nucleotide defects **[25].** Thus, due to elevated ROS and malfunctioning BER and NER the amount of unrepaired oxidative lesions, such as 8-oxo-dG, would increase over time further burdening the cells with dysfunctional TLS. Consequently, apart from re-replicated DNA **[9],** the unrepaired nucleotides may represent an additional source of replication fork stalling, collapse and DNA DSBs **[9]**.

**Figure 2.**
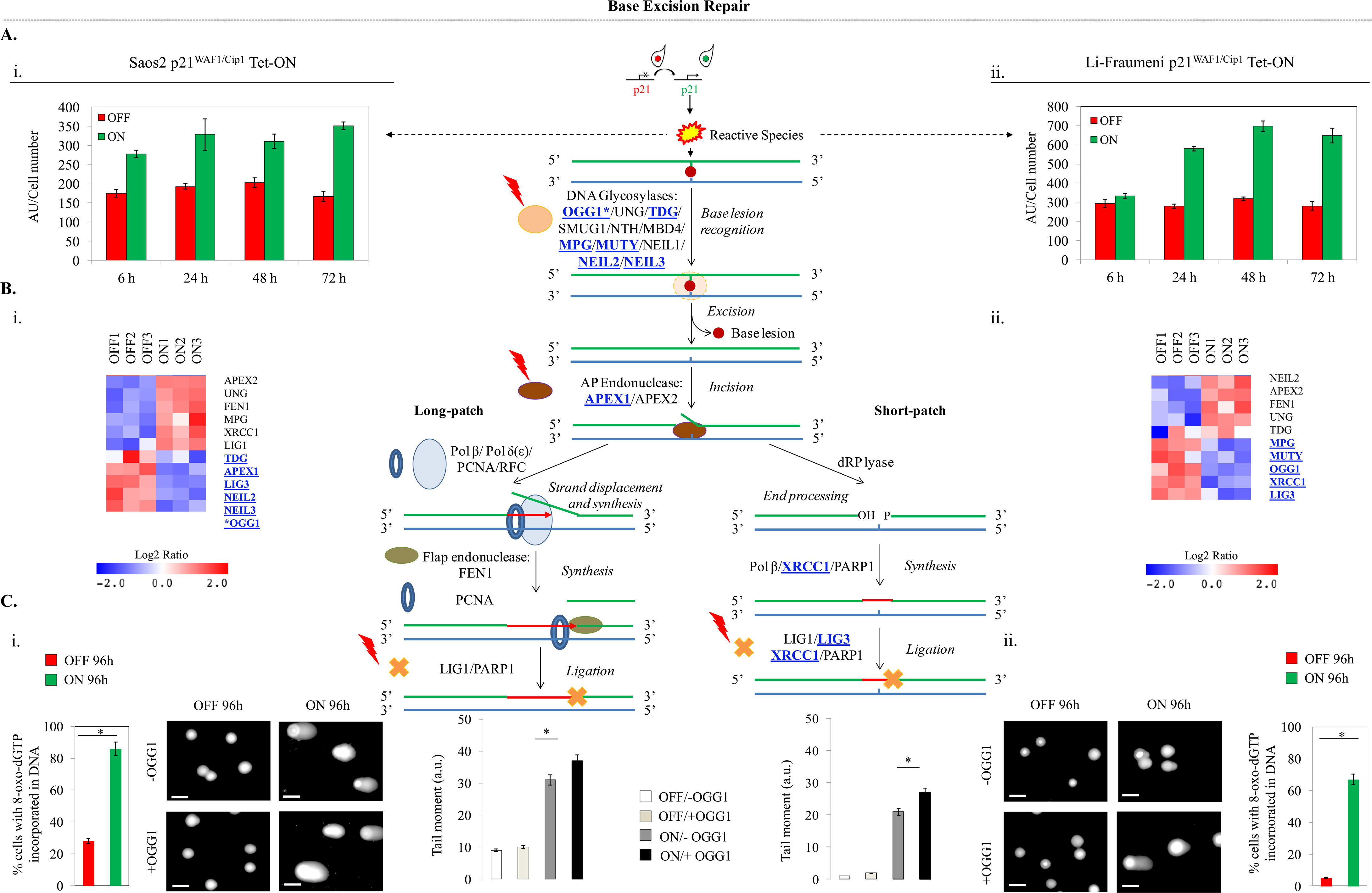
Decreased activity of the Base Excision Repair (BER) pathway in cells with sustained p21^WAF1/Cip1^ expression. (**ai-ii**) Increased Reactive Species (RS) levels were assessed with a DCFH-DA assay in (i) Saos2 and (ii) Li-Fraumeni cells with protracted p21^WAF1/Cip1^ expression [* *p*<0.05 (Saos2), * *p*=0.05 (Li-Fraumeni), *t*-test, error bars indicate SDs, n=3 experiments]. As shown in the middle panel RS production can lead to generation of nucleotide oxidative lesions. (**bi-ii**) RNAseq analysis showed that essential factors of the BER pathway were statistically significant down-regulated (*p*≤0.05) in 96h induced Saos2- and Li-Fraumeni-p21^WAF1/Cip1^ Tet-ON cells (see also **Suppl Fig 1** for specific real time RT-PCR validation). *Note that although in Saos2-p21^WAF1/Cip1^ Tet-ON cells OGG1 expression was not found by RNAseq analysis, specific real-time RT-PCR and microarray analysis (see also **Suppl Fig 1**) [**9**] confirmed its decreased expression. (**ci-ii**) Modified alkaline Comet assay demonstrated the presence of 8-oxoG in 96h induced Saos2- and Li-Fraumeni-p21^WAF1/Cip1^ Tet-ON cells, using 8-Oxoguanine glycosylase (OGG1) [* *p*<0.05 (Saos2), * *p*=0.05 (Li-Fraumeni), *t*-test, error bars indicate SDs, n=3 experiments]. Comet data were corroborated by an 8-oxo-dG specific assay measuring DNA incorporation of 8-oxo-dG in p21^WAF1/Cip1^ expressing cells [**24**], that is indicative of a lower OGG1 activity [* *p*<0.05 (Saos2), * *p*=0.05 (Li-Fraumeni), *t*-test, error bars indicate SDs, n=3 experiments]. Consequently, as depicted in the model of the middle panel, recognition and excision of the affected nucleotide lesion is impaired in the BER process. Middle panel depicts the components and steps during BER. BER pathway is responsible for removal of small lesions from DNA, especially oxidized, alkylated, deaminated bases and abasic sites. BER can be induced by oxidative stress and various genotoxic insults. Its specificity relies on the excision of base damage by glycosylases and AP endonucleases. In humans, the mechanism of BER involves the initial action of DNA glycosylases followed by the processing of the resulting abasic site either by the AP-lyase activity of the glycosylases or by the apurinic/apyrimidic endonucleases APE1/APE2 that incise the DNA strand. The resulting single-strand break can be processed by two BER subpathways. Either the short-patch branch is engaged, if a single nucleotide is replaced, or the long-patch branch, if 2-10 new nucleotides are synthesized. [OGG1: 8-oxoguanine DNA glycosylase; UNG: uracil DNA glycosylase; TDG: thymine DNA glycosylase; SMUG1: single-strand-selective monofunctional uracil-DNA glycosylase 1; NTH: DNA glycosylase and apyrimidinic (AP) lyase (endonuclease III); MBD4: methyl-CpG binding domain 4, DNA glycosylase; MPG: N-methylpurine DNA glycosylase; MUTY: adenine DNA glycosylase; NEIL1/2/3: Nei like DNA glycosylase 1/ 2/ 3; APEX1/2: apurinic/apyrimidinic endodeoxyribonuclease 1/2; POLB/POLD: DNA polymerase beta/ delta; PCNA: Proliferating cell nuclear antigen; RFC: replication Factor C; FEN1: Flap structure-specific endonuclease 1; LIG1/LIG3: DNA ligase 1/3; PARP1: poly(ADP-ribose) polymerase 1; XRCC1: X-ray repair cross complementing 1]

**Figure 3.**
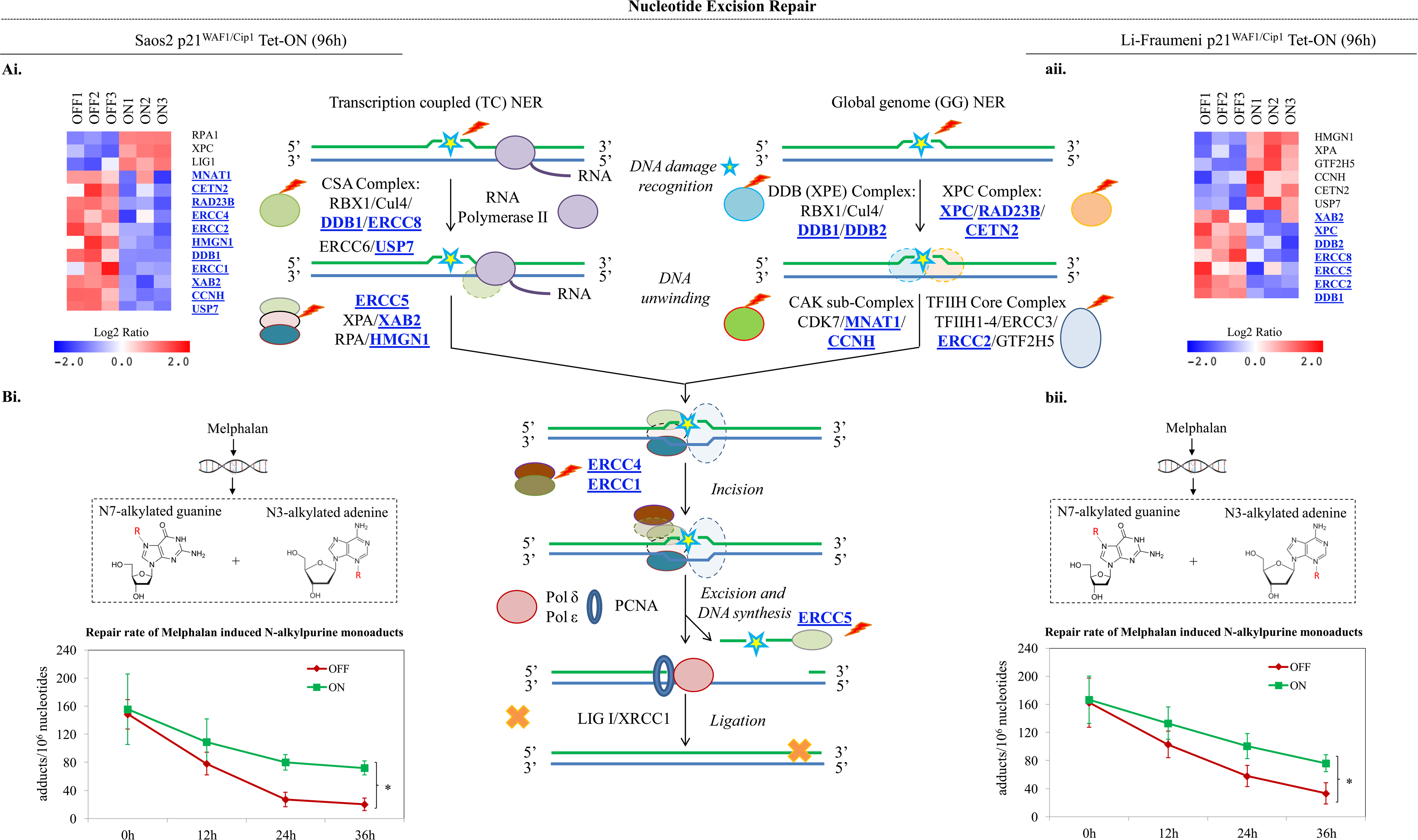
Decreased activity of the Nucleotide Excision Repair (NER) pathway in cells with prolonged p21^WAF1/Cip1^ expression. (**ai-ii**) RNAseq analysis showed that essential factors of the NER pathway were statistically significant down-regulated (*p* ≤0.05) (see also **Suppl Fig 1** for specific real-time RT-PCR validation) in 96h induced Saos2- and Li-Fraumeni-p21^WAF1/Cip1^ Tet-ON cells. (**bi-ii**) Decreased repair capacity of N-alkylpurine monoaducts in induced Saos2- and Li-Fraumeni-p21^WAF1/Cip1^ Tet-ON cells and treated with Melphalan [(4-(bis{2-chloroethyl} amino)-l-phenylalanine]. Middle panel depicts the components and function of NER. NER pathway is responsible for repair of bulky lesions, especially UV induced thymine dimers and 6,4-photoproducts, as well as non-bulky ones. Following DNA damage recognition, a short single-stranded DNA fragment that contains the lesion is removed. The remaining undamaged single-stranded DNA segment is used by DNA polymerase as a template to synthesize the complementary sequence. Final ligation to complete NER and formation of a double stranded DNA is carried out by DNA ligase. Depending on how the DNA damage is recognized, NER can be divided into two subpathways: transcription coupled NER (TC-NER) and global genome NER (GG-NER). While the two subpathways differ in how they recognize DNA damage, they share the same process for lesion incision, repair, and ligation. [RBX1: Ring-box 1; Cul4: Cullin 4; DDB1/2: Damage specific DNA binding protein 1/2; ERCC8 (CSA): ERCC excision repair 8, CSA ubiquitin ligase complex subunit; ERCC6 (CSB): ERCC excision repair 6, chromatin remodeling factor; USP7: Ubiquitin specific peptidase 7; ERCC5: ERCC excision repair 5, endonuclease; XPA: XPA, DNA damage recognition and repair factor; XAB2: XPA binding protein 2; RPA: replication protein A; HMGN1: high mobility group nucleosome binding domain 1; XPC: XPC complex subunit, DNA damage recognition and repair factor; RAD23B: RAD23 homolog B; CETN2: Centrin 2; CDK7: cyclin dependent kinase 7; MNAT1: MNAT1, CDK activating kinase assembly factor; CCNH: cyclin H; TFIIH1-4; Transcription/repair factor IIH 1-4; ERCC3: ERCC excision repair 3, TFIIH core complex helicase subunit; ERCC2: ERCC excision repair 2, TFIIH core complex helicase subunit; TTDA (GTF2H5/TFB5): general transcription factor IIH subunit 5]

### 3. Protracted p21^WAF1/Cip1^ expression fosters Rad52-dependent Break-induced Replication (BIR) and Single Strand Annealing (SSA)

Next we investigated whether mutational signatures could be used to identify potential repair pathway dysfunction. Notably, we found a complex pattern comprising concurrent signs of *signatures 6, 15* and *3* in the genomic landscape of the p21^WAF1/cip1^escaped cells **[8] (Fig. 4a,b)**. *Signature 6* is characterized by C→T substitutions at NpCpG sites and was reported to be strongly associated with inactivation of DNA mismatch repair genes **[8] (Fig. 4a),** a condition that was also observed in our settings **(Suppl. Fig 1b, d).** Although the origin of *signature 15* is still unknown, it is distinguished by C→T substitutions at GpCpN sites **[8] (Fig. 4a).** Interestingly, our transcriptome analyses showed reduced levels of BRCA2 and BRCA1 **(Fig. 4c),** a finding that could be relevant for *signature 3. Signature 3* is a mutational pattern characterized by deletions of up to 50-bp stretches of DNA with microhomologies at breakpoint junctions and inactivating mutations in *BRCA1* and *BRCA2.* Microhomologies are present in the majority of the novel breakpoints of the p21^WAF1/Cip1^escaped cells **[9],** and here we observed also down-regulation of both *BRCA1* and *BRCA2*, two critical components of homologous recombination repair **(Fig. 4c, Suppl Fig 2a)**. Concurrent loss of heterozygosity at the *BRCA2* locus was also noted **(Fig. 4c).** The low levels of BRCA2 along with the fact that Rad51 recombinase expression was reduced in the escaped cells **[9]** may leave Rad52 recombinase alone to drive Rad51-independent strand-annealing. Consistent with this notion, a strong endogenous Rad52 nuclear immunofluorescence signal was observed, in p21^WAF1/Cip1^induced cells (**Suppl Fig 3**), reflecting foci formation that co-localized with RPA **(Fig. 5a,b)**, suggesting a shift to Rad52-dependent recombination mechanisms **[9]**. In concordance with these findings we recorded a fast recruitment and maintenance of Rad52 at sites of DNA damage induced after UV irradiation ablation **(Fig. 5c; Suppl Video 2a-d)**.

**Figure 4.**
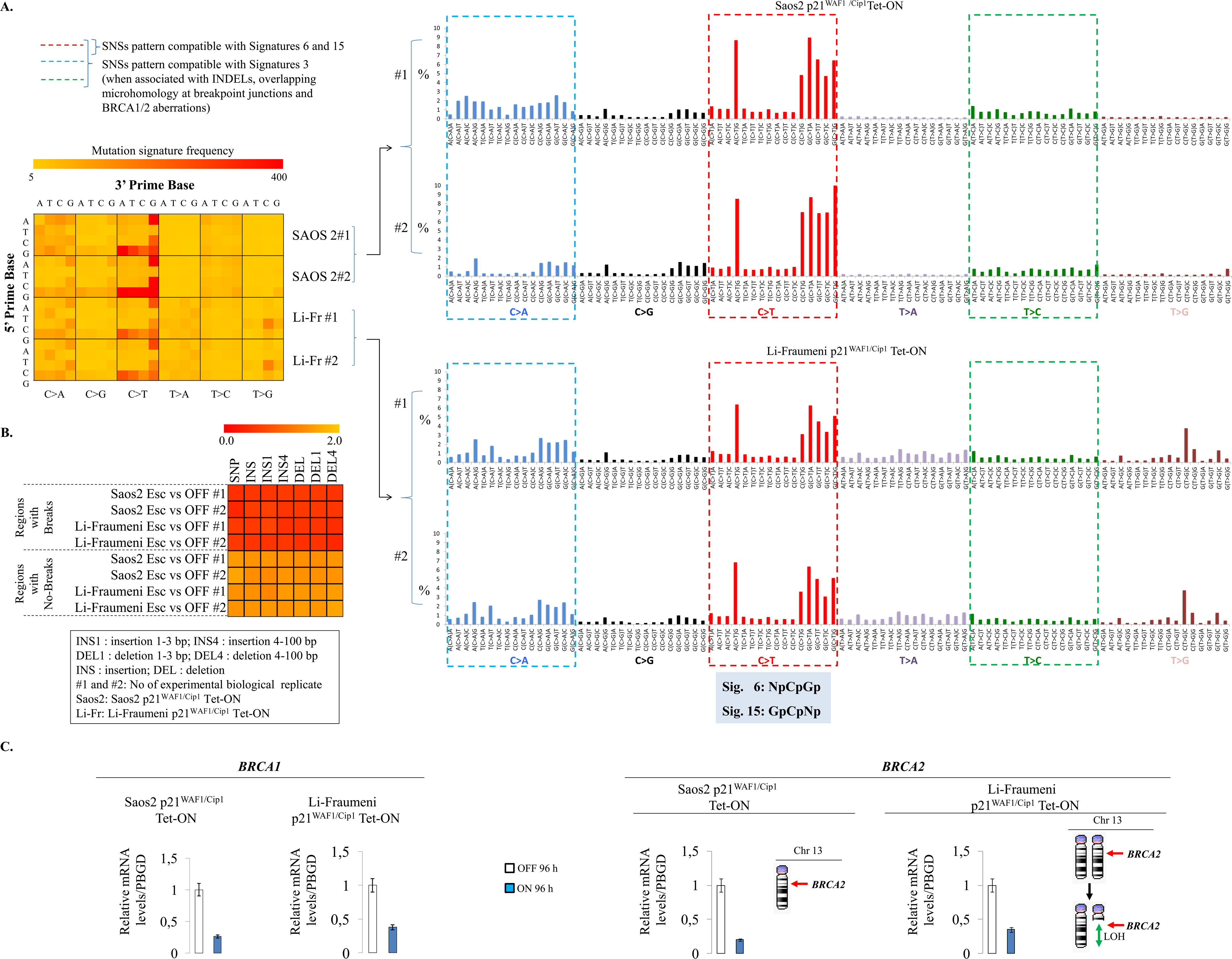
Extended p21^WAF1/Cip1^ over-expression shapes the mutational signature landscape. (a) Escaped (30 days induced) Saos2- and Li-Fraumeni-p21^WAF1/Cip1^ Tet-ON cells exhibit specific patterns of Single Nucleotide Substitution (SNSs). Heat-map showing the number of mutation-type at each mutation context, which was corrected for the frequency of each triplet in the human genome (hg19). Histograms present the mutation-type frequency at each mutation context from two biological replicates of escaped Saos2- and Li-Fraumeni-p21^WAF1/Cip1^ Tet-ON cells, respectively. Both presentations, show reproducible patterns of the mutational signatures 6, 15, 3 **[8]**. (**b**) Heat-map showing the association of SNSs, INS (nucleotide insertions) and DEL (nucleotide deletions) with the observed chromosomal breakpoints (± 50kb around the breakpoint) versus the remaining genome in escaped Saos2- and Li-Fraumeni-p21^WAF1/Cip1^ Tet-ON cells. (**c**) Real-time RT-PCR assessment of BRCA1 and BRCA2 mRNA expression in induced and non-induced Saos2 and Li-Fraumeni p21^WAF1/Cip1^ Tet-ON cells [* *p*<0.05 (Saos2), * *p*=0.05 (Li-Fraumeni), *t*-test, error bars indicate SDs, n=3 experiments]. Previous data [**9**] showed the monoallelic presence of chromosome 13 [that hosts the *BRCA2* locus (q13.10)] and loss of heterozygosity at the q arm in the Saos2- and Li-Fraumeni-p21^WAF1/Cip1^ Tet-ON cells, respectively.

**Figure 5.**
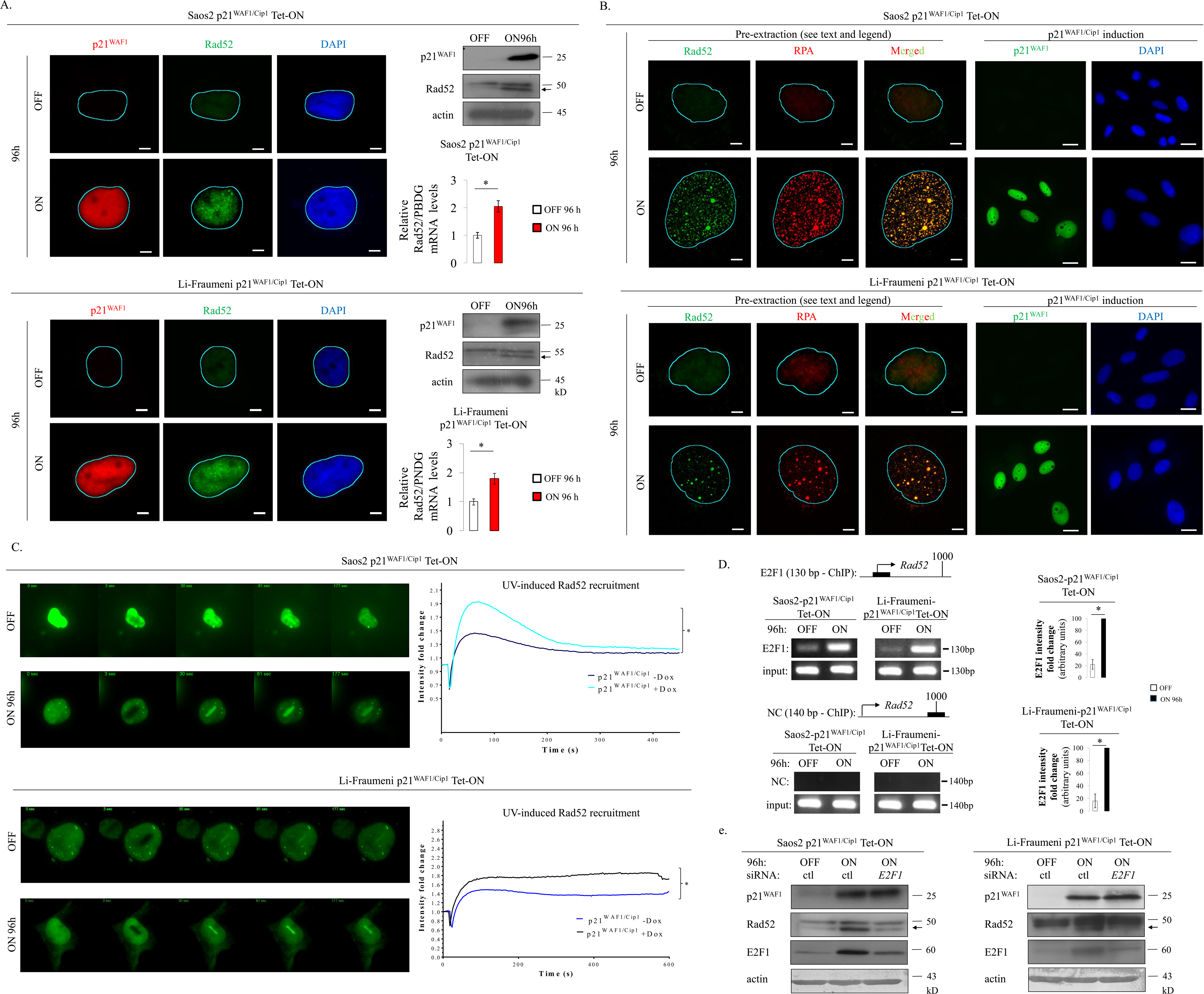
Rad52 increased expression, recruitment and foci formation at DSBs upon prolonged p21^WAF1/Cip1^ expression. (**a**) Immunofluorescent (IF) analysis showing increased Rad52 foci formation in 96h induced Saos2- and Li-Fraumeni-p21^WAF1/Cip1^ Tet-ON cells. Immunoblot (IB) and real-time RT-PCR assessment of Rad 52 expression levels in induced Saos2- and Li-Fraumeni-p21^WAF1/cip1^Tet-ON cells at the indicated time point [* *p*<0.05 (Saos2), * *p*=0.05 (Li-Fraumeni), *t*-test, error bars indicate SDs, n=3 experiments]. (**b**) IF analysis showing Rad52 and RPA foci formation and their co-localization in 96h induced Saos2- and Li-Fraumeni-p21^WAF1/cip1^Tet-ON cells. Saos2- and Li-Fraumeni-p21^WAF1/cip1^Tet-ON cells were pre-extracted with ice-cold PBS containing 0.2% Triton X-100 for 2 min on ice before fixation as previously described [**48**]. (**c**) IF confocal microscopy showing Rad52 loading on regions of damaged chromatin after UV-laser ablation in 96h induced Saos2- and Li-Fraumeni p21^WAF1/cip1^Tet-ON cells transfected with the YFP-Rad52 vector. Plots depict Rad52 recruitment kinetics at sites of DNA damage in the same cells, respectively. The average intensity of fluorescence at the site of damage and the total cell fluorescence in respect to time were quantified and plotted. Five cells in each condition of two independent experiments were processed. (**d**) *Rad52* promoter is occupied by E2F1 upon p21^WAF1/cip1^induction in Saos2- and Li-Fraumeni-p21^WAF1/cip1^Tet-ON cells, respectively, as assessed by chromatin immunoprecipitation (ChIP) (* *p*<0.05, *t*-test, error bars indicate SDs, n=3 experiments) (see also **Suppl Fig 4**). (**e**) Silencing of *E2F1* resulted in decreased Rad52 levels as assessed by immunoblot analysis in induced Saos2- and Li-Fraumeni-p21^WAF1/cip1^Tet-ON cells, respectively (* *p*<0.01, t-test, error bars indicate SDs, n=3 experiments). Actin serves as loading control; Ctl: control siRNA; h: hours, arrows indicate Rad52.

Intriguingly, the *RAD52* locus was almost deprived of SNSs and its protein expression was increased **(Fig 5a).** Notably, high protein levels were accompanied by increased transcription of Rad52, which is another interesting observation since most DNA repair factors are upregulated upon DNA damage, by post-transcriptional protein modifications that are faster compared to transcriptional control **[27].** To obtain a mechanistic insight into Rad52 transcriptional up-regulation, we conducted a bioinformatic analysis of its promoter. We found that the promoter sequence contains binding sites for several transcriptional factors including that for E2F1 (**Suppl. Fig. 4**). Notably, we previously showed that E2F1 is upregulated upon p21^WAF1/cip1^expression implying a putative functional link between E2F1 and Rad52 **[9].** The functionality of this potential signaling axis was supported by chromatin immunoprecipitation (ChIP) analysis of the *Rad52* promoter showing strong E2F1 binding in p21^WAF1/cip1^induced cells **(Fig. 5d).** Indeed, silencing of *E2F1* led to reduced *Rad52* expression, further supporting the above scenario **(Fig. 5e).**

In yeast, Rad52 is considered the lynchpin of homologous recombination by facilitating loading of Rad51 on single-stranded DNA (ssDNA) formed through DSB end resection; then Rad51-coated DNA invades the sister chromatid searching for homologous sequences forming D-loop structures **(Suppl. Fig. 2) [28-30]**. However in mammals, even though Rad52 retains strand-annealing activity **[31, 32],** Rad51 loading is mediated primarily by BRCA2 **[33-35],** implying that Rad52 may act as a back-up mechanism. This may explain why organismal development is unaffected in *RAD52*^*-/-*^ mice **[36, 37].**

Subsequently, analyzing the distribution of SNSs guided us in understanding which RAD52-dependent repair process took place in the p21^WAF1/cip1^-induced cells. Particularly, although at the genome-wide level the amount of SNSs was reduced in the escaped cells, the SNSs were interestingly clustered at the flanking regions in a number of novel breakpoint junctions **(Fig. 6a-c; Suppl Table 2).** This pattern of SNS clustering is termed “kataegis” **(Fig. 6a,b)** and it requires extensive tracts of single ssDNA that act as a substrate for cytidine deamination **(Suppl. Fig. 2a)**. The later is mediated by the APOBEC family of enzymes leading to C·G→T·A transitions and/or C·G→G·C transversions **[7, 38].** Break-induced replication (BIR), a homologous recombination (HR)-based repair route that allows replication re-start from collapsed replication forks **[39],** forms such long ssDNA tracts, representing a candidate to repair the p21^WAF1/cip1^induced DSBs **[9, 40].** During BIR a D-loop is formed followed by a replication fork at the one-ended DNA DSB. The D-loop is not dissolved, but moves together with the fork (migrating bubble) **[41, 42],** while DNA replication takes place in a conservative manner **[43] (Suppl. Fig. 2a).** In yeast, Rad52 seems to play a role in BIR **[44],** but its role in mammals has just started being elucidated [**30, 45, 46**]. Nevertheless, not all breakpoints in the escaped cells were flanked by clustered SNSs **(Fig. 6c,d; Suppl Table 2)** implying that more than one repair pathway is likely involved in processing the p21^WAF1/cip1^induced DSBs. Since a high frequency of microhomologies was observed in all novel breakpoints **[9]** we reasoned that a putative alternative repair route could be single strand annealing (SSA). SSA mediates annealing between two ssDNA ends containing homologous or microhomologous repeats, whereas the 3′-overhanging ends of the processed DSBs are trimmed by XPF-ERCC1 endonuclease **[47]** depriving the APOBEC editing enzymes from a single strand substrate for cytidine deamination **(Suppl. Fig. 2b).** Recently, it was shown that cells deficient in BRCA1 and 53BP1 relied on Rad52-dependent SSA to survive **[48].**

**Figure 6.**
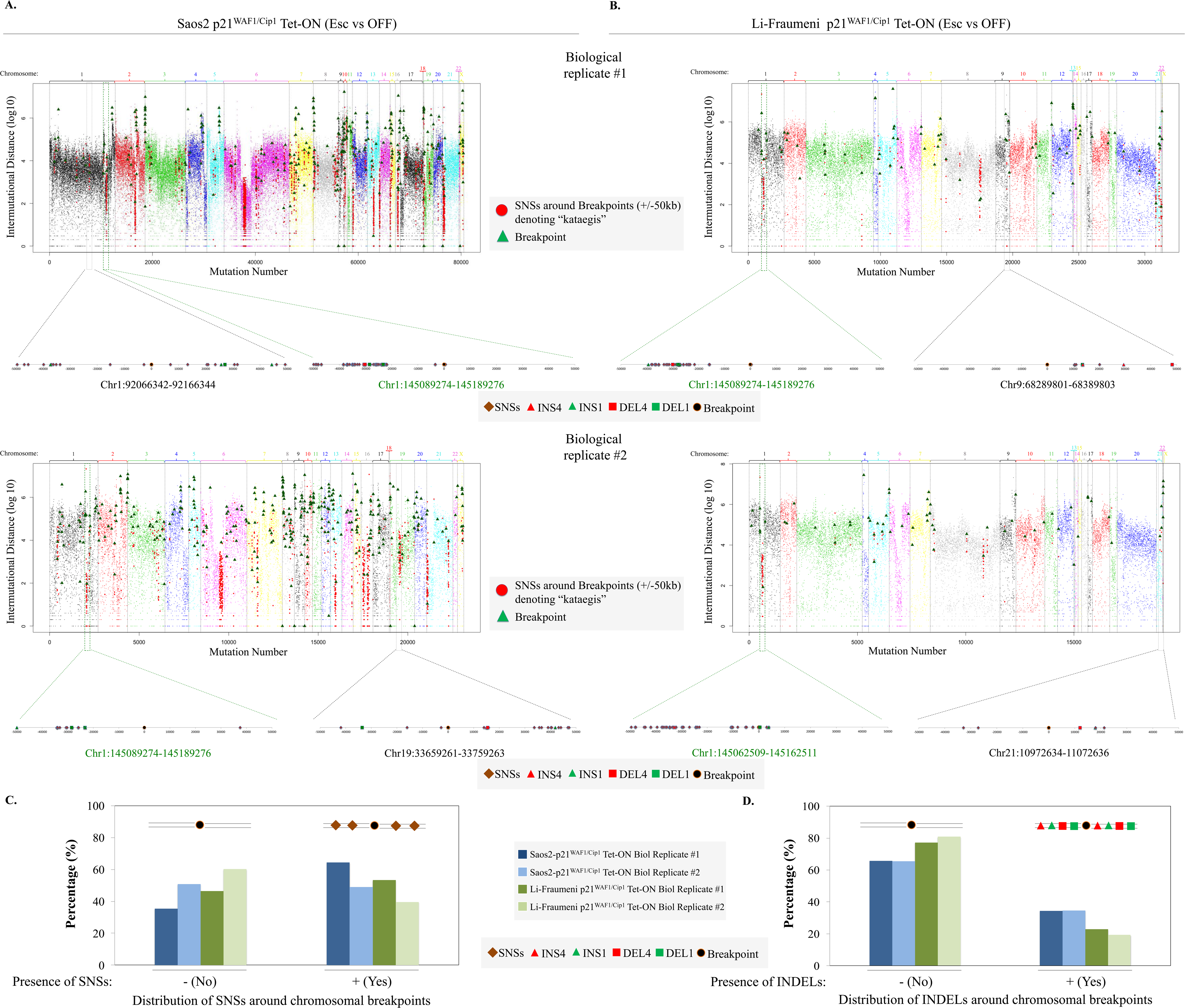
Single Nucleotide Substitutions (SNSs) cluster around chromosomal breakpoints in cells with continuous p21^WAF1/cip1^expression. (**a-b**) Diagrams depict a dense distribution of SNSs (red circles) in genome areas surrounding chromosomal breakpoints (green triangles), suggestive of kataegis phenomenon (small vertical arrows on top of diagrams), relative to disparate distribution of SNSs in the remaining genome of escaped (30 days induced), which are noted with different color for each chromosome, (a) Saos2- and (b) Li-Fraumeni-p21^WAF1/cip1^Tet-ON cells. Note that the total number of chromosomal breakpoints (green triangles) depicted is double as each side of a break corresponds to a different chromosomal arm (see **Suppl Table 2**). Dashed genome areas are depicted as magnifications of representative breakpoints. Green colored dashed areas signify representative breakpoints that show a high positional conservation in all experimental (biological) repetitions. Position of breaks and distribution of SNSs, INS (nucleotide insertions) and DEL (nucleotide deletions) in these examples is depicted in the corresponding subchromosomal magnifications. (**c-d**) Histograms depict the clustering frequency of (c) SNSs as well as (d) INS and DEL over all breakpoints in the genome of escaped (30 days induced) Saos2- and Li-Fraumeni-p21^WAF1/cip1^Tet-ON cells (see also **Suppl Table 2**).

To test the above assumption we monitored DSB repair using GFP-reporters, in which DSBs were generated by the nuclease I-SceI. The GFP-reporters were stably expressed in the Saos2 and Li-Fraumeni p21^WAF1/cip1^inducible systems, and were genetically modified in a way to monitor the main homology-dependent DSB repair modes: gene conversion (GC) and particularly synthesis-dependent strand annealing (SDSA), BIR and SSA **[39, 49] (Fig. 7).** DSB formation by I-SceI was followed by protracted p21^WAF1/cip1^expression for a period of 4 days. We observed that after day 4 the cells showed stronger fluorescence signals from the GFP-reporters monitoring BIR and SSA compared to the control p21^WAF1/cip1^-OFF cells, contrasting with the reduced fluorescence read-out from the cells expressing the GFP-SDSA reporter **(Fig. 7)**. The latter result was also consistent with the decreased expression of key components of the SDSA repair route seen after p21^WAF1/cip1^expression **(Fig. 4c, Suppl Fig 2a)**. Notably, exploiting the models of inducible p21^PCNA^ mutant that cannot interact with PCNA we noticed that BIR- and SSA-driven repair remained unaffected, confirming that the altered repair pattern observed in cells with induced wild-type p21^WAF1/cip1^was dependent on cellular effects mediated by the interplay of p21^WAF1/cip1^with PCNA during DNA replication **(Fig. 7)**. Consistent with such an overall model was also the reduced 53BP1 loading at sites of UV-induced DNA damage (**Fig. 1e**). 53BP1 fosters homology-directed DNA repair fidelity, but its level in cells is rate limiting, and its exhaustion signifies a shift to the error-prone SSA mechanism [**48**]. Collectively, these findings support our working hypothesis, and given the low levels of Rad51 and BRCA2, the data furthermore demonstrates the inability of SDSA to deal with p21^WAF1/cip1^triggered DSBs. To test the cause-effect relationship, we silenced *Rad52* to examine the dependency of BIR and SSA on Rad52, predicted by our model. Indeed, depletion of Rad52 led to a significant suppression of GFP fluorescence readouts from p21^WAF1/cip1^induced cells harboring both the BIR- and SSA-GFP repair reporters, thereby further supporting the requirement for Rad52 in these repair pathways in our experimental settings **[45, 48]**.

**Figure 7.**
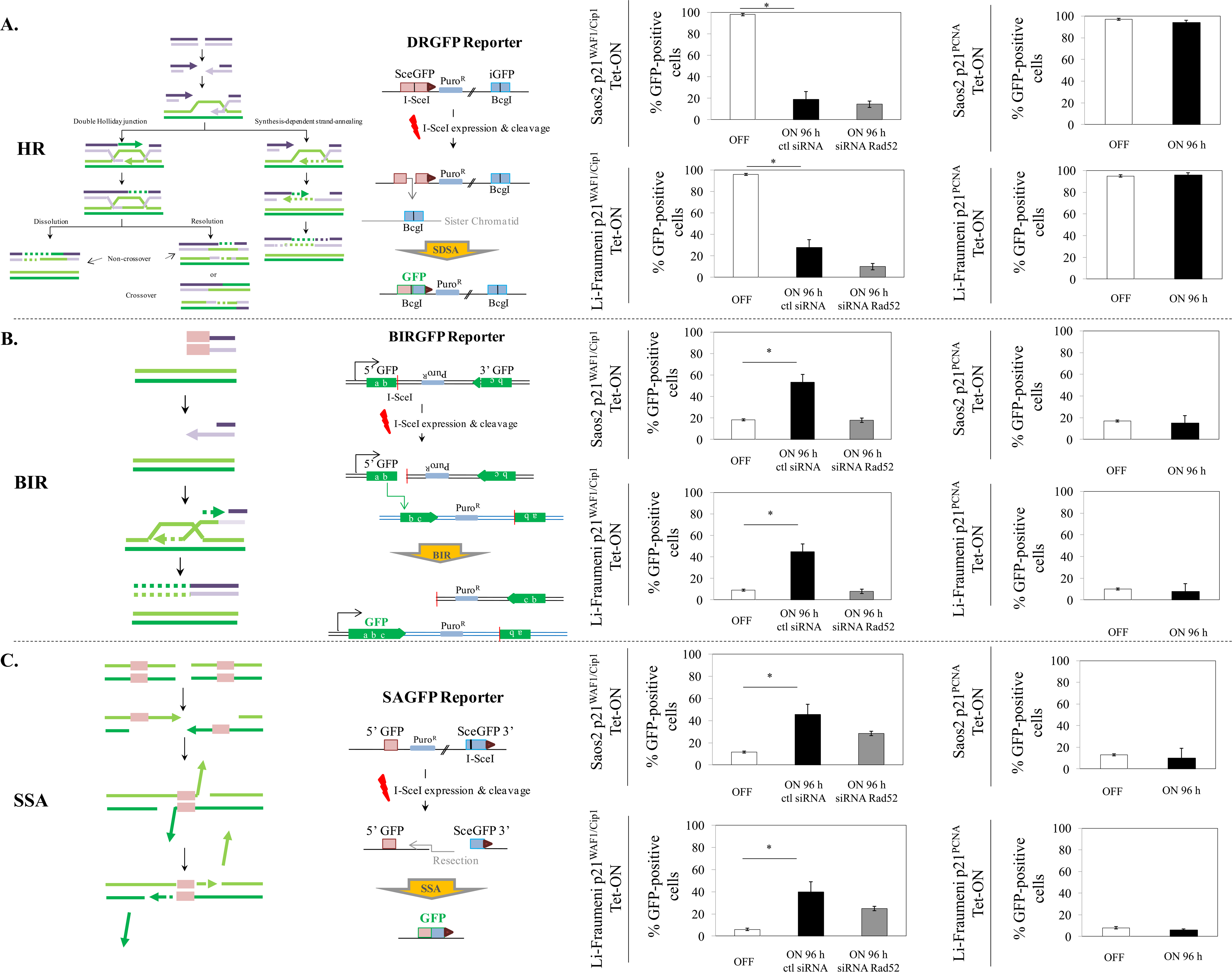
Prolonged p21^WAF1/cip1^expression promotes Rad52 dependent break induced-replication (BIR) and single-strand annealing (SSA) repair of DNA double strand breaks (DSBs). (**a**) Reduced synthesis-dependent strand annealing (SDSA) in 96h induced Saos2- and Li-Fraumeni-p21^WAF1/cip1^Tet-ON cells. Flow cytometry analysis (FACS) after p21^WAF1/cip1^induction in cells stably expressing a DR-GFP report vector and following I-SceI induced DSBs shows decreased SDSA activity (* *p*<0.05, *t*-test, error bars indicate SDs, n=3 experiments), regardless of Rad52 silencing. Similar manipulations in Saos2- and Li-Fraumeni-p21^PCNA^ Tet-ON cells showed no differences in SDSA in these cells. (**b**) Increased BIR activity in 96h induced Saos2- and Li-Fraumeni-p21^WAF1/cip1^Tet-ON cells. FACS after p21^WAF1/cip1^induction in cells stably expressing a BIR-GFP report vector and following I-SceI induced DSBs shows increased BIR activity (* *p*<0.05, *t*-test, error bars indicate SDs, n=3 experiments) that is suppressed upon Rad52 silencing. Similar experiment in Saos2- and Li-Fraumeni-p21^PCNA^ Tet-ON cells showed no effect on BIR function in these cells. (**c**) Increased SSA activity in 96h induced Saos2- and Li-Fraumeni-p21^WAF1/cip1^Tet-ON cells. FACS after p21^WAF1/cip1^induction in cells stably expressing an SA-GFP report vector and following I-SceI induced DSBs shows increased SSA activity (* *p*<0.05, *t*-test, error bars indicate SDs, n=3 experiments) that is dependent on Rad52. Similar experiment in Saos2- and Li-Fraumeni-p21^PCNA^ Tet-ON cells showed no effect on SSA function in these cells.

## Discussion

Genome maintenance is a fundamental prerequisite for preserving cellular life, both under physiological and pathological conditions. Even the highly unstable aberrant cancer genomes must be maintained within certain limits of genomic integrity, beyond which cells would die. Better understanding of the molecular mechanisms that allow genome destabilization that fuels cancer development and progression yet protect the cancer genome from too severe, fatal instability, is vital for the development of new therapeutic strategies **[6]**. Replication stress (RS) driven-genomic instability emerges as a major force promoting cancer evolution **[4, 50, 51].** To identify the DNA repair pathways that help cancer cells adapt and survive under such chronic stress is key to both, understanding tumorigenesis and finding clinically exploitable vulnerabilities of tumor cells.

Based on the results obtained in this study, we propose a concept whereby chronic expression of p21^WAF1/cip1^induces a dramatic rewiring of the cellular DNA repair pathway choices, providing further means in conjunction with deregulated replication to fuel genomic instability. Using human cellular models with inducible expression of p21^WAF1/cip1^that evokes replication stress [**9**], we now show that such scenario leads to suppression of the TLS-mediated repair, with ensuing defective processing of single nucleotide lesions, eventually resulting in replication fork stalling and collapse, generating DNA DSBs. Having in mind that most human cancers harbor p53 and p16^INK4^/pRb alterations [**52**], p53-independent expression of p21^WAF1/cip1^is one of the few remaining cellular guardians against the accumulating (pro)tumorigenic insults. Under such circumstances, p21^WAF1/cip1^-induced cellular senescence is often reversible and during this temporary anti-tumor response dramatic chromosomal remodeling takes place that favors over time the birth of aggressive off-springs **[9].** Here we provide mechanistic insights into the error-prone repair process that occurs during this genome-destabilizing evolutionary trajectory, by demonstrating that p21^WAF1/cip1^induced DSBs are commonly repaired by Rad52-dependent break-induced replication (BIR) and single strand annealing (SSA), as the synthesis-dependent strand annealing (SDSA) repair route was defected **(Fig. 4c, 7a; Suppl Fig. 2a)**. BIR is a highly error-prone homologous recombination (HR)-response that has been implicated in formation of high-frequency tandem segmental duplications found in cancer **[7, 39]**. Moreover, BIR introduces mutations in the newly synthesized DNA strand at a much higher rate than under conditions of conventional replication **[40, 53].** Likewise, SSA is associated with extensive DNA resection contributing to genome rearrangements and oncogenic transformation **[54]**.

Furthermore, we show that Rad52 was upregulated transcriptionally in a manner dependent on E2F1, a transcription factor that also drives G1/S transition and which is frequently overexpressed in cancer **[55]**. As the RB pathway is almost universally deregulated in tumors, E2F1 is often free from the restraining binding to pRB and hence capable of inducing Rad52. This, in turn, compensates for the reduced levels of Rad51 which is in short supply under stressful conditions **[56].** Transcriptional regulation, under genotoxic stress, as seen here for Rad52 was unexpected since most DNA damage response and repair proteins become rapidly upregulated via post-translational modifications that slow down the protein turnover **[27]**. Given the low levels of pivotal components (Rad51, BRCA1 and BRCA2) of the SDSA repair machinery **(Fig. 4c, Suppl Fig. 2a)**, the p21^WAF1/cip1^expressing cells resort to transcriptional upregulation of Rad52, a fact that reflects the increased repair needs required to cope with the p21^WAF1/cip1^-driven replication stress **[57]**. On the other hand, post-translational modifications of the chromatin bound fraction of Rad52 cannot be excluded, as recently reported **[45]**, and could further contribute to increased abundance of Rad52.

In contrast to yeast, where Rad52 represents a major factor in the first line of genome maintenance **[29],** in higher eukaryotes and mammals Rad52 seems to serve as a reserve player that can substitute for other repair options when those are compromised. Overall, this concept highlights Rad52 as a potential therapeutic target in tumors with inactive BRCA2 and helps explain why *Rad52* gene amplifications are selected for in human cancers **[58-60] (Suppl. Fig. 5).** Thus, targeting Rad52 could turn out to be a new way to therapeutically exploit vulnerabilities that occur selectively in cancer cells. Consistent with this idea, depletion of *Rad52* confers synthetic lethality in *BRCA2* deficient cells **[61, 62].**

## Conclusions

On the whole, the current study broadens our understanding of how chronic p53-independent p21^WAF1/cip1^expression, seen in a sizeable fraction of advanced human tumors **[9]** impacts the global DNA repair landscape and undermines genomic stability. The salient features of our model are the following: i) saturation of the CRL4^CDT2^ligase complex by p21^WAF1/cip1^impairs the turn-over of the replication licensing factors leading to their unscheduled accumulation, causing ii) genome re-replication and replication stress, as recently described **[9],** while iii) concurrent suppression of the DNA damage tolerance (TLS) pathway reduces the repair rate of the nucleotide lesions in an environment with dysfunctional error-free excision repair mechanisms of BER, NER and MMR. As a consequence of such grossly rewired DNA repair pathway choice, the rate of unrepaired nucleotide lesions increases, further raising the burden on the already limited TLS. Both features of DNA re-replication-induced replication stress and deficient TLS lead to enhanced formation of the highly deleterious DSBs that are repaired in an error-prone manner, by Rad52-dependent BIR and SSA, thereby fueling genomic instability and promoting cancer development **(Fig. 8).** We hope the concept proposed here may not only inspire further mechanistic studies, but also attempts to target Rad52 in cancer, as a way to selectively induce lethal chromosomal instability in Rad52-dependent cancers, while sparing normal tissues whose genome maintenance does not depend on Rad52. Lastly, the present study underscores the significance of identifying mutational signatures as they can unveil the repair procedure(s) that fuel genomic instability and thus highlight potential therapeutic targets for cancer treatment.

**Figure 8.**
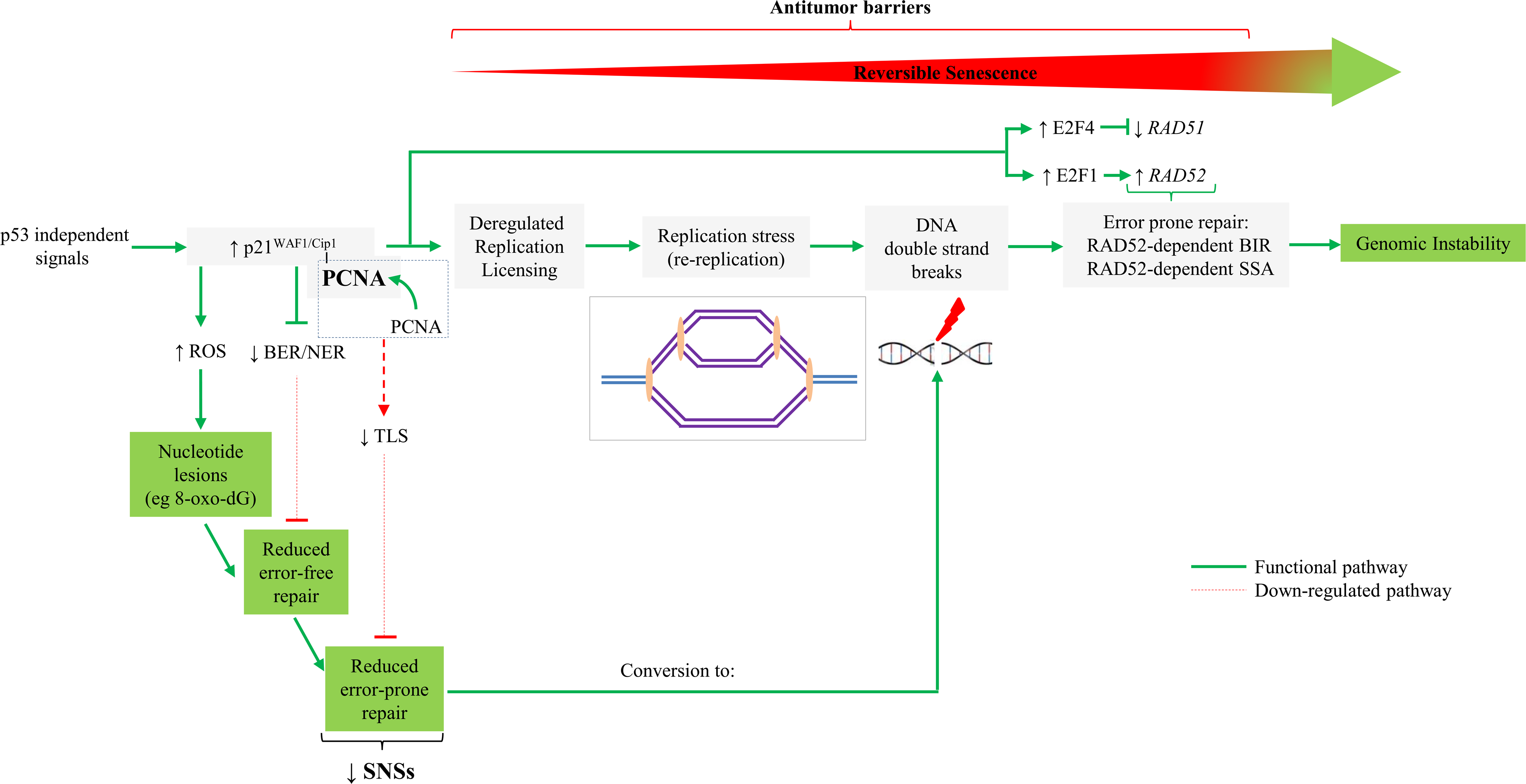
Proposed model depicting how p53-independent p21^WAF1/cip1^expression fuels Rad52–dependent error-prone double strand break repair promoting genomic instability. Sustained p53-independent p21^WAF1/cip1^induction leads to increased levels of nucleotide lesions mediated by elevated reactive oxygen species (ROS). Given the negative impact exerted by p21^WAF1/cip1^on the error free nucleotide repair mechanisms (BER and NER) a significant proportion of such base lesions escape unrepaired. This creates an additional repair “load” to the error prone repair mechanism of TLS which is further compromised by p21^WAF1/cip1^overexpression, leading to a decreased SNS load and in favor of DSBs. In turn, this further increases the DSB burden generated also through re-replication [**9**]. As components of SDSA are down-regulated, a shift to a Rad52-mediated error prone DNA repair takes place by invoking the BIR and SSA repair routes, fueling genomic instability. This repair switch is mediated by a shift in the balance between Rad51 and Rad52 levels as the former is suppressed by E2F4 [**9**] and the latter is induced by E2F1 (present study). DSBs: DNA double strand breaks; BER: base excision reapir; NER: nucleotide excision repair; TLS: translesion DNA synthesis and repair; SDSA: synthesis-dependent strand annealing; BIR: break induced repair; SSA: single strand annealing.

## Materials and Methods

### Cell lines and culture conditions

Inducible p21^WAF1/cip1^Tet-ON cell lines: Saos2-p21^WAF1/cip1^Tet-ON and Li Fraumeni-p21^WAF1/cip1^Tet-ON were maintained in High Glucose DMEM (Biosera) supplemented with 10% Tet System Approved FBS (Clontech) and 100 μg/ml penicillin and streptomycin (Invitrogen) and incubated at 37^°^C and 5% CO_2_. Induction of p21^WAF1/cip1^was conducted by treatment of the cell culture with 1 μg/ml doxocycline (Applichem) [**9**].

### siRNA and vector transfections

Rad52 (Thermo Scientific) siRNA gene silencing was performed as previously described, following the manufacturer’s instructions [**9**]. YFP-RAD52 [**48**] and Pol-κ vectors were transfected as previously described [**63**].

### Protein extraction, cell fractionation and immunoblotting

Protein extraction and cell fractionation was performed as described before [**9**]. Thirty micrograms of protein from total extracts per sample were adjusted with Laemmli buffer (Sigma) and loaded on acrylamide/bis-acrylamide gels. Gel electrophoresis, transfer to PVDF membrane (Millipore) and signal development with nitro blue tetrazolium/5-bromo-4-chloro-3-indolylphosphate (NBT/BCIP) solution (Molecular Probes) or chemiluminescence were performed as previously described [**9**]. Alkaline phosphatase-conjugated anti-mouse or anti-rabbit as well as Horse Radish Peroxidase conjugated anti-mouse, anti-rabbit and anti-sheep secondary antibodies (1:1000 dilution) (Cell Signaling) were used.

Primary antibodies utilized were: anti-p21^WAF1/cip1^(mouse, Santa Cruz, sc-6246, 1:400 for IB), anti-RAD52 (mouse, Santa Cruz, sc-365341, 1:100 for IF), anti-β-actin (rabbit, Cell Signaling Tech, 4967s, 1:1000 for IB), anti-PCNA (mouse, Cell Signaling Tech, 2586s, 1:1000 for IB), anti-Ubiquityl-PCNA (rabbit, Cell Signaling Tech, 13439s, 1:1000 for IB), anti-polη (rabbit, Santa Cruz, sc5592, 1:200 for IB), anti-β-tubulin (rabbit, Abcam, ab6046, 1:1000 for IB), anti-lamin B1 (rabbit, Abcam, ab16048, 1:1000 for IB).

### Indirect Immunofluorescence

Indirect immunofluorescence analysis was performed as previously published **[9**]. Regarding identification of RAD52 foci parameters of the aforementioned process have been set as indicated by Ochs and collaborators [**48**] and by Sotiriou and collaborators [**45**]. For all RPA and Rad52 IF co-localization experiments (**Fig. 5b**) Saos2- and Li-Fraumeni-p21^WAF1/cip1^Tet-ON cells were pre-extracted with ice-cold PBS containing 0.2% Triton X-100 for 2 min on ice before fixation as previously described [**48**]. Secondary antibodies were Alexa Fluor 488 donkey anti-sheep (Abcam, ab150177, 1:500) and Alexa Fluor 568 goat anti-mouse (Invitrogen, no. A11031, 1:500). Image acquisition of multiple random fields was automated on **a DM 6000 CFS Upright Microscope (Confocal Leica TCS SP5 II), or** a ScanR screening station (Olympus) and analyzed with ScanR (Olympus) software, or a Zeiss Axiolab fluorescence microscope equipped with a Zeiss Axiocam MRm camera and Achroplan objectives, while image acquisition was performed with AxioVision software 4.7.1. Primary antibodies utilized were: anti-p21^WAF1/cip1^(mouse, Santa Cruz, sc-6246, 1:200 for IF), anti-RAD52 (sheep, [**48]**, 1:100 for IF and mouse, Santa Cruz, sc-365341, 1:100 for IF).

### Image acquisition

Live DNA damage protein recruitment kinetics were observed using an Olympus IX83 inverted microscope system. DNA damage irradiation was induced on the same system using a coupled UVA (355nm) pulsed laser (teemphotonics PNV-M02510) and a theoretical pulse duration of less than 350psec. Subnuclear irradiations were performed on a 8μm linear ROI with the use of a total of 60-90 pulses subdivided in 3 repeats, with 2.3% of the total laser power. Pulse irradiation calibration was titrated through γH2Ax post damage spatial organisation on MCF7 cancer cells as previously described [**64**]. For time-lapse acquisition an Olympus Apochromat 63x/ 1.2NA water immersion lens, a Hamamatsu ORCA Flash 4.0. sCMOS camera system were used. The microscope was equipped with a temperature/humidity and CO2 incubation system (CellVivo) and with a 6-LED system (Lumencor) as light source.

Brief powerful laser ablation inscribed cell location within the glass volume of the coverslip below the cells of interest. This technique enables easy location of the marked field of views on any microscope under transmission contrast.

Z-stack imaging was conducted on a Leica SP5 TCS equipped with Hybrid Detector and a 60x/ 1.4NA oil immersion lens. A z-step of 0.72um was used and a total of 18 stacks were obtained per nucleus.

#### Cell culture

For live-cell experiments, cancer cells were plated on Ibidi glass bottom dishes (idibi μ-dish 35mm 81156) in phenol red-free, Minimum Essential Medium Eagle (MEM). L-Glutamine 2mM, Hepes 25mM final concentration and 10% Fetal Bovine Serum (FBS) were added.

#### Image analysis

Kinetics analyses were quantified using Fiji distribution of ImageJ (v 2.0.0-rc-30/1.49s). Images were background corrected by subtracting the mean intensity value of an area outside the cell (ROI3). Values of corrected mean intensity of the site of damage (RO1’) and the corrected total fluorescent-area mean intensity were obtained at each time point (ROI2’). Cells were normalised for the different protein expression levels and acquisition photobleaching with the use of this formula:

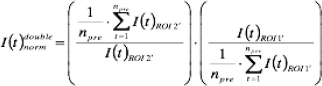

### Chromatin immunoprecipitation (ChIP) assay

ChIP assay was performed as previously described [**63**]. A 130bp fragment in the *Rad52* promoter and a 140bp amplicon, located approximately 1000bp from the transcription start site **(Suppl Fig. 4b)**, were amplified. Primers and annealing temperatures are provided in **Suppl Table 3**. PCR reactions containing 1% of the total chromatin extract used in the immunoprecipitation reactions were used as inputs.

### Mammalian translesion synthesis (TLS) assay

Cells were co-transfected with a plasmid mixture containing the gap-lesion plasmid (kan^R^), a control gapped plasmid without a lesion (cm^R^), and the carrier plasmid pUC18 (amp^R^). After allowing time for gap filling and lesion bypass, plasmids were extracted using alkali, such that only filled in plasmids remained intact. To assay the fraction of filled in plasmids, the plasmid mixture was transformed into an indicator *E. coli recA* strain and plated in parallel, on LB-kan plates (to select for plasmids that underwent TLS) and LB-cm plates (to select for the control filled in plasmid GP20-*cm*). TLS in this case was calculated by the ratio of kan^R^/cm^R^ *E. coli* transformants. Specifically, the cells were co-transfected with a DNA mixture containing 50 ng of a gap-lesion plasmid (GP-BPG1; kan^R^), 50 ng of a gapped plasmid without lesion (GP20-*cm*, cm^R^), and 10 ng of the carrier plasmid pUC18, using jetPEI/DNA complexes. The percentage of lesion bypass gap filling was calculated by dividing the number of GP-BPG1 transformants (number of colonies on LB-kan plates) by the number of corresponding GP20-cm transformants (number of colonies on LB-cm plates). When desired, plasmids were extracted from kan^R^ colonies, and the sequence opposite the lesion was determined by DNA sequence analysis **[18]**.

### TLS assay in E. coli

Gapped plasmids carrying the BP-G adduct (BP-BPG1; kan^R^) and the control plasmid without the adduct (GP20; cm^R^) were used in parallel to transform UV-irradiated *E. coli* cells. The cells were UV irradiated at 20 Jm^−2^, followed by a recovery period of 30 min at 37^°^ C, after which they were transformed in parallel with the gapped plasmids with and without the BP-G adduct using the Ca-MOPS method. Survival was calculated by dividing the number of transformants obtained with the gap-lesion plasmid by the number of transformants obtained with the gapped plasmid without the lesion. In this assay, results from two parallel transformations were compared. Although no internal control was included, the same stock of competent cells was used for the gapped plasmids with and without the lesion, and the reliability of the results was assured by performing multiple experiments for each pair of gapped plasmid constructs **[18]**.

### Comet assay

In order to determine the endogenous (background) levels of oxidatively-induced DNA damage we performed the sensitive technique of Single Cell Gel Electrophoresis (SCGE; Comet assay) under alkaline (denaturing) conditions as previously described [**63**]. We also used as damage probe the E. coli repair enzyme OGG1 (New England Biolabs) to detect the specific presence of 8-oxo-dGuanine (8-oxo-dG) [**24**]. An increase in the tail moment (TM) suggests higher levels of DNA damage. Cells were observed under a Zeiss Axiolab fluorescence microscope equipped with a monochrome CCD camera. Analysis was conducted with Cometscore software (Tritek). All experiments were performed in triplicates.

### Melphalan Assay

Melphalan [(4-(bis{2-chloroethyl} amino)-l-phenylalanine] belongs to the nitrogen mustard class of chemotherapeutic agents, used in the treatment of certain haematological maligancies. Its mode of action is by alkylating the DNA, generating predominantly N-alkylpurine monoaducts and to a minor extent interstrand cross-links (ICLs), the formation of which are dependent on these monoaducts. These lesions primarily affect the N-7 position of guanines and to a lesser degree the N-3 position of adenines. These monoaducts are almost exclusively repaired by the Nucleotide Excision Repair (NER) mechanism.

To measure the induction and repair of melphalan-derived DNA adducts, a specific assay is applied [**26**]. In principle, extracted and digested DNA from treated cells undergoes depurination of N-alkylated bases and conversion of these sites to single strand breaks. Following hybridization with probes for various genes, at consecutive time points after drug treatment, quantitative data of obtained signals for the analyzed genes is correspondingly estimated. Average frequency of N-alkylpurines in the fragments of interest are calculated from the fraction of DNA in the corresponding gene band from the treated sample as compared to that from untreated control sample.

### 8-oxo-dG assay

A previously described assay based on the property of avidin to bind with high specificity to 8-oxo-dG was used for the 8-oxoG measurements [**24**]. Briefly, cells were fixed in methanol at -20^°^C for 20 min and incubated for 15 min in TBS, 0.1% Triton X-100. Blocking was performed in 15% FBS, 0.1% Triton X-100 in TBS for 2 h at room temperature (RT). Cells were then incubated with 10 μg/ml Alexa488-conjugated avidin (Invitrogen) in blocking solution for 1 h at 37^°^ C. Next they were rinsed twice in TBS, 0.1% Triton X-100 for 5 min each round at RT. After a quick rinse in distilled water, DNA was counterstained with ToPro3-Iodide (LifeTechnologies) for 15 min at RT, followed by a final rinse in TBS.

Coverslips were mounted with ProLongGold (Invitrogen) and cells were observed under a Zeiss Axiolab fluorescence microscope equipped with a monochrome CCD camera. Analysis was conducted with NIH-imageJ, with respect to mean intensity in the nucleus (To-Pro3 served as a DNA reference).

### Measurement of Intracellular Levels of Reactive Oxygen Species

Intracellular levels of reactive oxygen species (ROS) were calculated using the DCFH-DA assay. In details, the cells were plated in a 96-well plate at a density of 10,000 cells/well and when approx. 80% confluent they were treated with doxocycline (1 μg/ml) in DMEM supplemented with 10% (v/v) FBS until confluence. At the indicated time points DCFH-DA (10 μM) was added and after a further incubation of 1 hour measurements (excitation wavelength: 480 nm, emission wavelength: 530 nm) were taken in a FLUOStar OPTIMA microplate reader (BMG Labtech GmbH, Ortenberg, Germany) using the MARS Data Analysis Software). After this measurement, the number of cells was estimated and ROS levels were expressed as fluorescence units per cell number [**65**].

### cDNA preparation and real-time quantitative PCR with reverse transcription (rt_RT-qPCR)

cDNA generation and rt_RT-qPCR analysis was performed as described before [**9**]. The reaction was performed in a StepOne Real time machine (Life Technologies) using Universal MasterMix II without UNG containing SYBR (Life Technologies) and 200nM primers. Signal analysis was carried out using the StepOne v2.3 software. Primers and annealing temperatures are provided in **Suppl Table 3**.

### DR-GFP, SA-GFP and BIR-GFP reporter assays

Saos2-p21^WAF1/cip1^Tet-ON cells harboring the GFP based reporter constructs for synthesis-dependent strand annealing (DR-GFP), single strand annealing (SA-GFP) and break induced replication (BIR-GFP), were generated by transfection with these DSB repair reporters, followed by selection of stably transfected clones [**39, 49**]. To monitor the repair of an I-SceI-generated DSB, cells were transiently transfected with 1μg of the I-SceI expression vector HA-ISceID44A (Addgene #59424) with effectene transfection reagent (Qiagen). DSB repair efficiency, upon induction of p21^WAF1/cip1^, was determined by quantifying GFP-positive cells via flow cytometry FACS Calibur (Becton Dickinson), 48h after transfection.

### High-throughput whole-genome analyses

#### Whole-genome sequencing (WGS) analysis

WGS library preparation, alignment and breakpoint identification was performed as described before [**9**]. Samtools mpileup and vcftools [**66**] were used for identification and filtering of the SNPs and INDEls. SNPs and INDELs that were unique in the “escaped” cells were normalized based on the depth of the sequencing for each experiment.

#### RNA-seq analysis

RNA was collected from non-induced and 96h induced (4 d) Saos2 p21 Tet-ON cells, as well as from non-induced and 96h induced (4 d) Li-Fraumeni p21 Tet-ON cells (three biological replicates for each condition). RNA-seq library preparation and analysis of 75bp paired-end reads procedure was performed in the Greek Genome Center (GGC) of Biomedical Research Foundation of Academy of Athens (BRFAA). Tophat2 (2.0.9) [**67**] was used for data alignment with the use of “- - sensitive” option to the hg19 genome version, while HT-seq count algorithm [**68**] was used for assigning aligned reads to the human transcriptome. Identification of the differentially expressed genes was performed with R/Bioconductor and DESeq [**69**] algorithm and genes with absolute fold-change≥1.5 and p-value≤0.05 were considered as differentially expressed between induced and non-induced cells. RNAseq data are submitted under the accession number SRAXXXX.

## List of Abbreviations

TLS: Translesion DNA Synthesis;
BER: Base Excision Repair;
NER: Nucleotide Excision Repair;
MMR: Mismatch Repair;
SNSs: Single Nucleotide Substitutions;
DSBs: Double Strand Breaks;
RS: Replication Stress,
BIR: Break Induced Replication;
SSA: Single Strand Annealing;
SDSA: Synthesis Dependent Strand Annealing;
DDT: DNA Damage Tolerance;
PRR: Post Replication Repair;
HR: Homologous Recombination;
GC: Gene Conversion;
ROS: Reactive Oxygen Species;
AP sites: apurinic/apyrimidinic sites;
BPDE: Benzo[a]pyrene Diol Epoxide;
8-oxo-dG: 8-oxo-dGuanine;
OGG1: 8-Oxoguanine glycosylase

## Declarations

### Ethics approval and consent to participate

Not applicable

### Consent for publication

All authors are aware of the content and agree with the submission.

### Availability of data and material

RNAseq data are submitted under the accession number SRAXXXX.

### Author contributions

PG, GP, IS, ISP and CG: cell culture and manipulations, siRNA/plasmid-transfections, immunoblots, ChIP, 8-oxo-dG assay, IF, and DNA repair assays assays. AK: RNA extraction, RT-PCR, figures preparation. AG: Comet assay. NNG and ZL: live image analyses and interpretation. AP: WGS, RNA-seq and bioinformatic analysis. VS: Melphalan Assay. USNG and ZL: TLS assays and analyses. CL and JL: RPA and Rad52 IF co-localization experiments and provided Rad52 in-house specific antibody. JL, LS, CSS, and JB: data analysis and interpretation, assistance in manuscript preparation. VGG: experimental design, guidance, manuscript preparation and writing.

#### Acknowledgements

We would like to thank Prof TD Halazonetis for providing the constructs for synthesis-dependent strand annealing (DR-GFP), single strand annealing (SA-GFP) and break induced replication (BIR-GFP); Dr E Soutoglou for kindly donating the I-SceI expression vector HA-ISceID44A and Chr Zampetidis for expert help.

## Supplementary Figure legends

**Suppl Figure 1. Reduced expression of components of the Base Excision Repair (BER), Nucleotide Excision Repair (NER) and Mismatch Repair (MMR) pathways in cells with sustained p21^WAF1/cip1^expression.** (**a, c**) Expression status of components of the BER, NER and MMR pathways as assessed by real-time RT-PCR in 96h induced Saos2- and Li-Fraumeni-p21^WAF1/cip1^Tet-ON cells, validating the high-throughput expression results (see also **Figs. 2, 3**) (*p* < 0.01, t-test, error bars indicate SDs, n=3 experiments). Note that for OGG1 expression status, data from previous microarray RNA analysis in Saos2-p21^PCNA^ Tet-ON cells was also available [**9**]. (**b, d**) RNAseq analysis of essential factors of the MMR pathway in 96h induced Saos2- and Li-Fraumeni-p21^WAF1/cip1^Tet-ON cells. The lower right panels depict the components and steps implicated during MMR. The MMR pathway is responsible for recognition and repair of mismatched bases generated due to misincorporation during DNA replication. In eukaryotes the mechanism of MMR involves the initial recognition of the mismatched base by MutSα/β, followed by incision from the endonuclease activity of MutLα of the 3’- or 5’-side of the mismatched base on the discontinuous strand. Subsequently, EXO I exonuclease excises the resulting DNA segment, in cooperation with the single-stranded DNA-binding protein RPA. The resected DNA strand is resynthesized by DNA polymerase δ and DNA Ligase. [MSH2/3/6: MutS homolog 2/3/6; MLH1/3: MutL homolog 1/3; PMS1/2: PMS1/2 homolog 1, mismatch repair system component; EXO1: Exonuclease 1]

**Suppl Figure 2. Models depicting proposed function of synthesis-dependent strand annealing (SDSA), break-induced replication (BIR) and single-strand annealing (SSA) repair mechanisms.** (**aii**) In the SDSA model, double-stranded breaks (DSB) are repaired by homologous recombination, without formation of a double Holliday junction, producing non-crossover products. In this model, following DSB recognition (MRN complex: MRE11/RAD59/NBS) the DSB ends are recessed (by CtIP and EXO I, facilitated by SLX1/SLX4). The generated single-stranded 3' overhangs are then coated with the RPA protein, while RAD51 loading is facilitated (BRCA2/PALB2/DSS1). Subsequently, strand invasion by Rad51 single-stranded DNA nucleoprotein filaments takes place, forming a D-loop. The invading 3’ strand extends along the recipient homologous DNA duplex by DNA polymerase (Pol δ) in a 5’ to 3’ direction. As a result the D-loop “moves” in this process, which is termed as bubble migration DNA synthesis. The single Holliday junction also slides down the DNA duplex in the same direction in a process called branch migration, displacing the extended strand from the template strand. The newly synthesized 3' end of the invading strand is then able to anneal to the other 3' overhang in the damaged chromosome through complementary base pairing. After the strands anneal, a small flap of DNA can sometimes remain. Such flaps are removed, and the process finishes with the resealing, also known as ligation (LIG), of any remaining single-stranded gaps. [**47, 53, 54, 71-73**] (**aii**) The break-induced repair (BIR) pathway is a homology-dependent repair route of one sided DSBs. During DNA replication, DSBs arising from stalled or collapse replication fork as well as DSBs emerging from erosion of uncapped telomeres are one end sided. They are considered to be repaired by strand invasion into a homologous duplex DNA followed by replication to the chromosome end (conservative replication). Specifically, the DNA at the DSB end is 5’ to 3’ recessed (by CtIP and EXO I facilitated by SLX1/SLX4), generating single-stranded 3' overhangs that are coated with the RPA protein, while RAD51 loading is facilitated (BRCA2/PALB2/DSS1). Subsequently, strand invasion by Rad51 single-stranded DNA nucleoprotein filaments takes place, forming a D-loop. The D-loop is not dissolved but moves together with the replication fork (migrating bubble - arrow showing direction of movement). The D-loop progresses with concomitant leading and lagging strand DNA synthesis (Pol D3). As lagging strand DNA synthesis delays, long single strand stretches (red arrow denoted) are produced. Following strand displacement gaps are filed and ligation (LIG) restores continuity of the DNA strand. The BIR pathway is considered mutagenic since it results in loss of heterozygosity or in a nonreciprocal translocation, if an ectopically template is employed. [**47, 53, 54, 71-73**] (b) The single-strand annealing (SSA) pathway is a homology-dependent repair route of DSBs occurring between two repeat sequences. In contrast to the SDSA pathway, the SSA mechanism does not rely on the use of a separate homologous DNA strand for recombination. Instead, the SSA route requires only a single DNA duplex, using the repeat sequences flanking the DSB, as the homologous recombination sequences needed for repair. The steps of the SSA pathway are as follows. Initially, upon damage recognition (MRN complex: MRE11/RAD59/NBS) the DNA around the DSB is recessed (CtIP and EXO I facilitated by SLX1/SLX4), generating single-stranded 3' overhangs that are coated with the RPA protein, thus preventing these overhangs from sticking to themselves. Subsequently, Rad52 binds the repeat sequences on each side of the break, and aligns them to enable the two complementary repeat sequences to anneal. Subsequently, the 3' non-homologous flaps are excised by the XPF/ERCC1 nucleases (denoted by black arrows), whose recruitment is facilitated by MSH2/MSH3. Gaps are filed and ligation (LIG) restores continuity of the DNA strands. The SSA pathway is considered mutagenic since the DNA sequence between the repeats and one of the two repeats are lost. [**47, 53, 54, 71-73**]

**Suppl Figure 3. Increased Rad52 expression upon sustained p21^WAF1/cip1^expression.** Quantitative analysis of Rad52 IF signal intensity in Saos2-p21^WAF1/cip1^Tet-ON cells, showing the high levels of Rad52 expression upon p21^WAF1/cip1^induction. A.U.: arbitrary units.

**Suppl Figure 4. E2F1 mediates *Rad52* transcriptional up-regulation upon extended p21^WAF1/cip1^expression.** Subchromosomal localization, gene organization and promoter sequence of the human *Rad52* locus. Analysis of the *Rad52* gene promoter (retrieved from Ensembl) using the online bioinformatic tool “GPMiner” (http://gpminer.mbc.nctu.edu.tw/index.php) that uses the Transfac database of mammalian transcriptional regulatory elements, revealed several putative transcription binding sites including E2F1.

## Supplementary Video legends

**Supplementary Video 1a.** Video time-laps imaging showing Polκ recruitment at sites of DNA damage after UV induced laser ablation in non-induced Saos2-p21^WAF1/cip1^(Tet-OFF) cells.

**Supplementary Video 1b.** Video time-laps imaging showing Polκ recruitment at sites of DNA damage after UV induced laser ablation in induced Saos2-p21^WAF1/cip1^(Tet-ON) cells.

**Supplementary Video 1c.** Video time-laps imaging showing Polκ recruitment at sites of DNA damage after UV induced laser ablation in non-induced Li-Fraumeni-p21^WAF1/cip1^(Tet-OFF) cells.

**Supplementary Video 1d.** Video time-laps imaging showing Polκ recruitment at sites of DNA damage after UV induced laser ablation in induced Li-Fraumeni-p21^WAF1/cip1^(Tet-ON) cells.

**Supplementary Video 2a.** Video time-laps imaging showing Rad52 recruitment at sites of DNA damage after UV induced laser ablation in non-induced Saos2-p21^WAF1/cip1^(Tet-OFF) cells.

**Supplementary Video 2b.** Video time-laps imaging showing Rad52 recruitment at sites of DNA damage after UV induced laser ablation in induced Saos2-p21^WAF1/cip1^(Tet-ON) cells.

**Supplementary Video 2c.** Video time-laps imaging showing Rad52 recruitment at sites of DNA damage after UV induced laser ablation in non-induced Li-Fraumeni-p21^WAF1/cip1^(Tet-OFF) cells.

**Supplementary Video 2d.** Video time-laps imaging showing Rad52 recruitment at sites of DNA damage after UV induced laser ablation in induced Li-Fraumeni-p21^WAF1/cip1^(Tet-ON) cells.

